# Closer appendage spacing augments metachronal swimming speed by promoting tip vortex interactions

**DOI:** 10.1101/2021.03.03.433771

**Authors:** Mitchell P. Ford, Arvind Santhanakrishnan

## Abstract

Numerous species of aquatic invertebrates, including crustaceans, swim by oscillating multiple closely spaced appendages. The coordinated, out-of-phase motion of these appendages, known as “metachronal paddling”, has been well-established to improve swimming performance relative to synchronous paddling. Invertebrates employing this propulsion strategy cover a wide range of body sizes and shapes, but the ratio of appendage spacing (*G*) to the appendage length (*L*) has been reported to lie in a comparatively narrow range of 0.2 < *G/L* ≤ 0.65. The functional role of *G/L* on metachronal swimming performance is unknown. We hypothesized that for a given Reynolds number and stroke amplitude, hydrodynamic interactions promoted by metachronal stroke kinematics with small *G/L* can increase forward swimming speed. We used a dynamically scaled self-propelling robot to comparatively examine swimming performance and wake development of metachronal and synchronous paddling under varying *G/L*, phase lag, and stroke amplitude. *G/L* was varied from 0.4 to 1.5, with the expectation that when *G/L* is large, there should be no performance difference between metachronal and synchronous paddling due to a lack of interaction between vortices that form on the appendages. Metachronal stroking at non-zero phase lag with *G/L* in the biological range produced faster swimming speeds than synchronous stroking. As *G/L* increased and as stroke amplitude decreased, the influence of phase lag on the swimming speed of the robot was reduced. For smaller *G/L*, vortex interactions between adjacent appendages generated a horizontally-oriented wake and increased momentum fluxes relative to larger *G/L*, which contributed to increasing swimming speed. We find that while metachronal motion augments swimming performance for closely spaced appendages (*G/L* < 1), moderately spaced appendages (1.0 ≤ *G/L* ≤ 1.5) can benefit from metachronal motion only when the stroke amplitude is large.

## 1. Introduction

Metachronal paddling is a fluid transport mechanism used in a variety of biological functions— including locomotion, feeding, and mucus transport—across a wide number of distantly related taxa (Campos et al. 2012, Catton et al. 2011, Sensenig et al. 2010, Sleigh and Barlow 1980, Sleigh et al. 1988, Wong et al. 1993, van Duren and Videler 2003). Among organisms that use metachronal paddling for locomotion, there are diverse body morphologies, ranging from microscopic paramecia with numerous cilia to macroscopic soft-bodied ctenophores and hard-bodied crustaceans. Crustaceans in particular consistute one of the most abundant macroscopic taxa on Earth, and themselves include a broad range of morphological diversity, from copepod nauplii with body lengths on the order of 10 μm (Lenz et al. 2015) to adult lobsters of body sizes exceeding 1 m. Free-swimming crustaceans use rhythmic oscillations of multiple closely spaced swimming appendages, where appendage geometry as well as number and location of swimming appendages vary across species. Most crustaceans typically possess between 3 to 8 pairs of thoracic and/or abdominal swimming appendages (Alexander 1988, Campos et al. 2012, Catton et al. 2011, Schabes and Hamner 1992, Van Duren and Videler 2003) which they stroke in series starting from the posterior to the anterior with a time delay (phase lag) between adjacent pairs. The spacing or gap between appendages (*G*) of metachronal swimmers has been reported to occupy a rather narrow range (Murphy et al. 2011), with the ratio of *G* to appendage length (*L*) ranging from as low as *G*/*L*=0.2 for some copepods to *G*/*L*=0.65 for Pacific krill. However, the effect of varying *G/L* on metachronal swimming performance and the value of *G/L* beyond which swimming performance becomes independent of *G/L* are currently unknown.

Little is known about the fluid-structure interactions occurring due to the coordinated motion of appendages at the scales at which crustaceans swim. Variations in body size and *G/L* can result in altering the fluid dynamic mechanisms responsible for thrust and drag generation by an animal, as the Reynolds number based on pleopod length, stroke frequency, and stroke amplitude ranges over several orders of magnitude among crustaceans, from 10^0^ to 10^4^ (Campos et al. 2012, Lim and DeMont 2009, Murphy et al. 2011, Schabes and Hamner 1992, Van Duren and Videler 2003). Previous studies have examined the wake (Catton et al. 2011, Lim and DeMont 2009, Murphy et al. 2011, Yen et al. 2003), stroke kinematics (Campos et al. 2012, Lim and DeMont 2009, Murphy et al. 2011, van Duren and Videler 2003), and swimming performance (Campos et al. 2012, Lenz et al. 2015, Murphy et al. 2011, Murphy et al. 2013, Schabes and Hamner 1992, van Duren and Videler 2003) in several crustacean species. There are several parameters that could influence swimming performance, including body and pleopod geometries, stroke frequency, stroke amplitude, phase lag, and stroke waveform. Previous experimental and numerical studies have shown that using metachronal paddling, when performed with closely spaced paddles, can result in increased swimming performance relative to synchronous paddling across a wide variety of system designs and Reynolds numbers (Alben et al. 2010, Ford et al 2019, Ford and Santhanakrishnan 2021, Granzier-Nakajima et al. 2020, Hayashi and Takagi 2020, Larson et al. 2014, Takagi 2015, Zhang et al. 2014). While it is plausible that placing paddles in close proximity of each other promotes constructive vortex interactions that benefit swimming performance, such hypotheses have not been tested previously in organism-level and modeling studies.

In this study, we use experiments on a self-propelling metachronal swimming robot to examine the effects of changing *G*/*L* on the swimming wake and mechanical performance. We vary the gap between paddles, the phase lag between the motion of adjacent paddles, and the stroke amplitude of angular paddle motion. High-speed videos of the robot swimming were used to determine the effects of changing these morphological and kinematic parameters on swimming performance. Flow visualization using particle image velocimetry (PIV) measurements was used to examine the hydrodynamic interactions that can explain observed changes in swimming performance.

## 2 Materials and Methods

### 2.1 Experimental setup

Using a programmable robotic paddling platform developed in a previous study (Ford and Santhanakrishnan 2021), but with a generalized flat plate body (**Figure 1**), we tested paddling motion of physical models varying in *G/L* from 0.4 to 1.5. Experiments were performed in a 244 cm long × 65 cm wide × 77 cm tall glass aquarium filled with 300 gallons of a solution of approximately 85% by volume of glycerin and 15% by volume of water (kinematic viscosity of fluid mixture, *v*=100 mm^2^ s^−1^, density *ρ*=1220 kg m^−1^). A 1 m long air bearing (model A-108.1000, PI (Physik Instrumente) L.P., Auburn, MA, USA) was mounted above the aquarium on a custom-built aluminum frame, which allowed for low-friction movement along the longitudinal axis of the robotic model.

**Figure 1.**
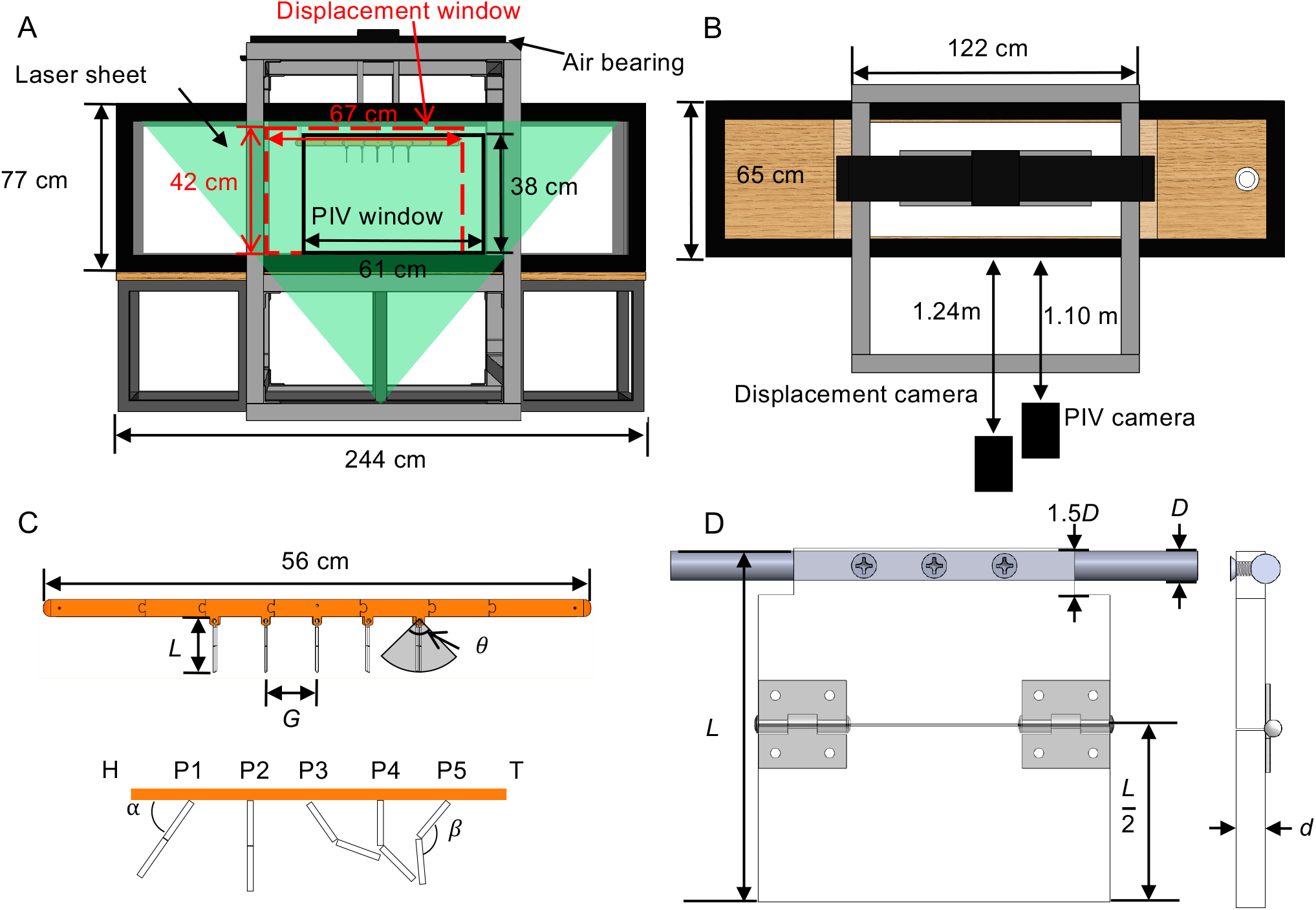
Schematic diagram of the experimental setup used in this study. (A) Front view of the aquarium used in this experiment. An air bearing (top, black) was used to allow near-frictionless motion along the longitudinal axis of the paddling robot. Two different camera windows were used for data collection.. The displacement window (red dashed lines) was centered towards the “head” of the robotic model prior to the start of the motion in order to assess motion throughout the travel. The PIV window (black solid lines) was centered towards the “tail” of the model, which allowed for the wake to be resolved throughout the motion. (B) Top view of the aquarium showing camera positions for displacement and PIV recordings. (C) Close-up views of the model indicating paddle length (*L*), gap between paddles (*G*), stroke amplitude (*θ*), appendage angle (*α*) and hinge angle (*β*), where H=head and T=tail of the model. Paddles are sequentially numbered as P1 to P5 as shown in the bottom close-up view, such that P1 and P5 are the anterior and posterior paddles, respectively. (D) Close-up view of the paddles. L=7.62 cm, with hinges located halfway down the paddle length. Paddle thickness, d=0.3 cm. Motion is driven by a 0.6 cm diameter aluminum shaft.

The robotic model was designed as a platform for comparative studies of swimming performance across different species and behaviors. Body segments were interchangeable to easily vary *G/L*. Paddles were 7.62 cm on each side with an aspect ratio (length divided by width) of 1. A passive hinge was mounted halfway down the paddle, which allowed the interior joint angle (*β* in **Figure 1**) of the paddle to rotate freely between approximately 180 degrees (during power stroke) and 100 degrees (during recovery).

Dynamic scaling was achieved by matching both the Reynolds number and the Strouhal number of paddling krill. We previously showed that by matching Reynolds number and kinematics of the robot with those of Antarctic krill, we were able to recover similar Strouhal numbers and swimming performance (Ford and Santhanakrishnan 2021). Reynolds number (*Re*) was defined based on the average tip velocity of the paddle (*U*_tip,mean_), the paddle length (*L*), and the *v* of the fluid medium in which the robotic model was submerged, as shown in the equation below:

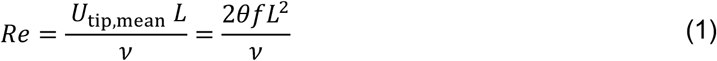

where *f* is the stroke frequency and *θ* is the stroke amplitude, and *U_tip,mean_* = 2*θfL*. While changing experimental variables *f, θ, L* and *v* result in changing *Re*, changing the phase lag and the gap between paddles (*G*) have no explicit effect on *Re*. Motion of each of the 5 paddles was independently controlled according to their respective stroke amplitudes and phase lags as described in the next subsection. In addition to constant *v* described earlier, stroke frequency (*f*) was maintained constant at 2.5 Hz for all tests conducted in this study.

### 2.2 Kinematics

A series of 5 stepper motors were used to drive paddle motion and were controlled using a custom LabVIEW program that prescribed angular positions of the paddles at 10 ms increments, as in Ford and Santhanakrishnan (2021). Angular resolution of the motors was 0.018 degrees/step. Upper appendage kinematics (*α* in **Figure 1**) were prescribed with stroke amplitude (*θ*) ranging from 55° to 95° in increments of 10°, and phase lag (*ϕ*) ranging from 0% to 20% of cycle time. To avoid collision of adjacent paddles during the stroke cycle, *θ* was limited to a maximum of 75° for *G*/*L*=0.5 and a maximum of 85° for *G*/*L*=1.0. Appendage angle (*α*) and hinge angle (*β*, denoted in **Figure 1**) were tracked from high-speed videos of the model during self-propulsion and were tracked in ImageJ software (National Institutes of Health, Bethesda, MD, USA). Examples of prescribed and tracked appendage angles for *G*/*L*=0.5, 1.0 and 1.5 are shown in supplementary material (**Figure S1, A-C**) for *ϕ*=20% and *θ*=75°. Achieved paddle motion matched closely with prescribed *α*. Tracked *β* profiles are shown for a representative test condition in the supplementary material (**Figure S1, D-F**). The paddles can be observed to fold-in during recovery stroke (*t/T*=0.5-1), as evidenced by decreasing *β* during this portion of the cycle.

### 2.3 Displacement

High-speed swimming videos were recorded using a Phantom Miro M110 camera (Vision Research, Wayne, NJ, USA) with sensor size of 25.6 × 16.0 mm and resolution of 1280 × 800 pixels. The camera was positioned 124 cm from the front of the tank. A 60 mm fixed focal length lens was mounted to the camera, with the aperture set to f/2.8. This provided a field of view 67 cm in length to record the displacement of the swimming model. Videos were recorded at 250 frames per second (100 frames per paddling cycle at *f*=2.5 Hz), and displacement was tracked using DLTdv7 (Hedrick 2008) in MATLAB (The MathWorks Inc, Natick, MA, USA). Swimming speed was calculated as the average speed over a stroke cycle. The model started from rest and reached a steady swimming speed after several stroke cycles had elapsed, and an example of the time-resolved displacement of a similar model can be found in our previous paper (Ford and Santhanakrishnan 2021). Equation 2, which defines the calculation for swimming speed, is shown below.

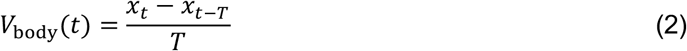

where *V*_body_ is the swimming speed, *x_t_* is displacement at time *t*, *x_t−T_* is displacement at the same phase from the previous cycle (*t − T*), and *T* is cycle time (*T*=1/*f*=400 ms).

### 2.4 Particle image velocimetry (PIV)

Two-dimensional time-resolved PIV measurements were performed to visualize the evolution of the paddling wake. The high-speed camera used in displacement measurements was also used for PIV, but moved slightly closer to the aquarium. The PIV camera was positioned 110 cm from the front surface of the aquarium to give a field of view 61 cm wide, as opposed to the 67 cm field of view used in the displacement tracking (**Figure 1A,B**). Illumination for the PIV recordings was provided by a 527 nm wavelength high-speed single-cavity Nd:YLF laser (Photonics Industries International, Ronkonkoma, NY, USA) with maximum pulse energy of 30 mJ/pulse at 1 kHz pulse frequency, and a maximum repetition rate of 10 kHz. PIV cross-correlation was performed in DaVis 8.4 (LaVision GmbH, Göttingen, Germany), using a two-pass cross-correlation with a first pass window of size 32 × 32 pixels and 50% overlap, and a second pass window of size 12 × 12 pixels and 50% overlap. From the PIV velocity fields, the out-of-plane vorticity component (*ω_z_*) was calculated according to the equation below:

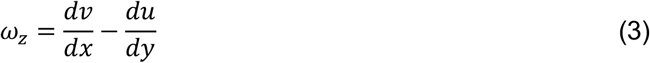

where *v* is the vertical component of velocity, *u* is the horizontal component of velocity, *x* is horizontal position in the flow field and *y* is vertical position in the flow field. *d/dx* and *d/dy* indicate infinitesimal derivatives with respect to *x* and *y* directions, respectively.

### 2.5 Momentum flux

Cycle-averaged momentum fluxes were calculated at specific locations in the PIV field of view to obtain estimates of cycle-averaged force (Ford et al. 2019). Horizontal momentum flux (HMF) per unit depth was calculated at several locations along the length of the robotic body, at measured locations between paddles P1 and P5 (paddle numbers are indicated in **Figure 1C**). Additionally, vertical momentum flux (VMF) per unit depth was calculated at various depths below the body, ranging from 0.5L to 3.5L below the tip of the fully extended paddle. Momentum flux per unit depth is defined as the integral of fluid momentum across a line. VMF and HMF were calculated according to the following equations:

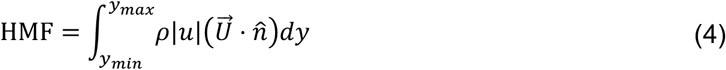

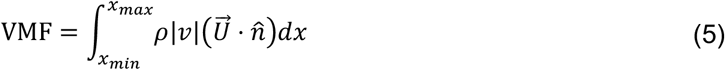

where 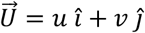 is the two-dimensional velocity vector at a particular location in the flow (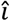 and 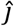 denote the unit normal vectors in *x* and *y* directions, respectively), *ρ* is the fluid density measured to be 1220 kg m^−3^, *y_min_* is the lowest position in the camera frame, *y_max_* is the highest position in the camera frame, *x_min_* is the leftmost position in the camera frame, *x_max_* is the rightmost position in the camera frame, and 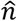 is the unit vector perpendicular to the direction of interest (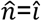 for HMF; 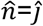 for VMF).

In addition to momentum flux, total momentum in a volume represents how much fluid is moved by the paddling motion. Total momentum is the product of mass and velocity within the fluid volume. In this case, momentum per unit depth is calculated as:

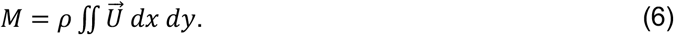

The box used for the momentum calculation did not cover the entire field of view of the camera, but instead covered the length of the robotic model (56 cm) and had a height of 30 cm. Horizontal and vertical components of momentum were compared to determine the angle of the wake. The wake angle was determined using the equation:

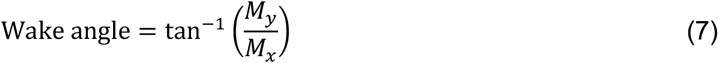

where *M_y_* and *M_x_* are the vertical and the horizontal components of the momentum *M*, respectively.

## 3 Results

### 3.1 Swimming speed

Swimming speed was calculated according to the definition given in **Equation 2**. After an initial period of acceleration, average swimming speed approached a steady value which was recorded as the steady swimming speed of the robot. The means and standard deviations of steady swimming speed across three independent trials for each condition are shown in **Figure 2**. In general, swimming speed decreases with increasing *G*/*L*, as well as increasing with increasing *θ*. The best performance (highest swimming speed) was a close match between *G/L* = 1.0 with *ϕ* = 10-15% and *θ* = 85° and *G/L* = 0.4 with *ϕ* = 10-15% and *θ* = 75°. For the largest appendage spacing with *G/L* =1.5, changing the phase lag *ϕ* showed no effect on steady swimming speed. Since the swimming speed does not change with varying phase lag at *G*/*L*=1.5, it can be inferred that the hydrodynamic interactions between paddles do not change. As *G/L*→∞, the wakes of individual paddles do not interact, and swimming speed becomes independent of *ϕ* and *G*/*L*. This is seen as increasing *G/L* from 1.0 to 1.5 has little effect on swimming speed (particularly at low values of *θ*) and varying *ϕ* has almost no effect on swimming speed at *G*/*L*=1.5. Regardless of *θ*, changing *ϕ* has the least effect on swimming speed at *G*/*L*=1.5, and the most effect for *G/L* ranging between 0.4 to 0.6.

**Figure 2.**
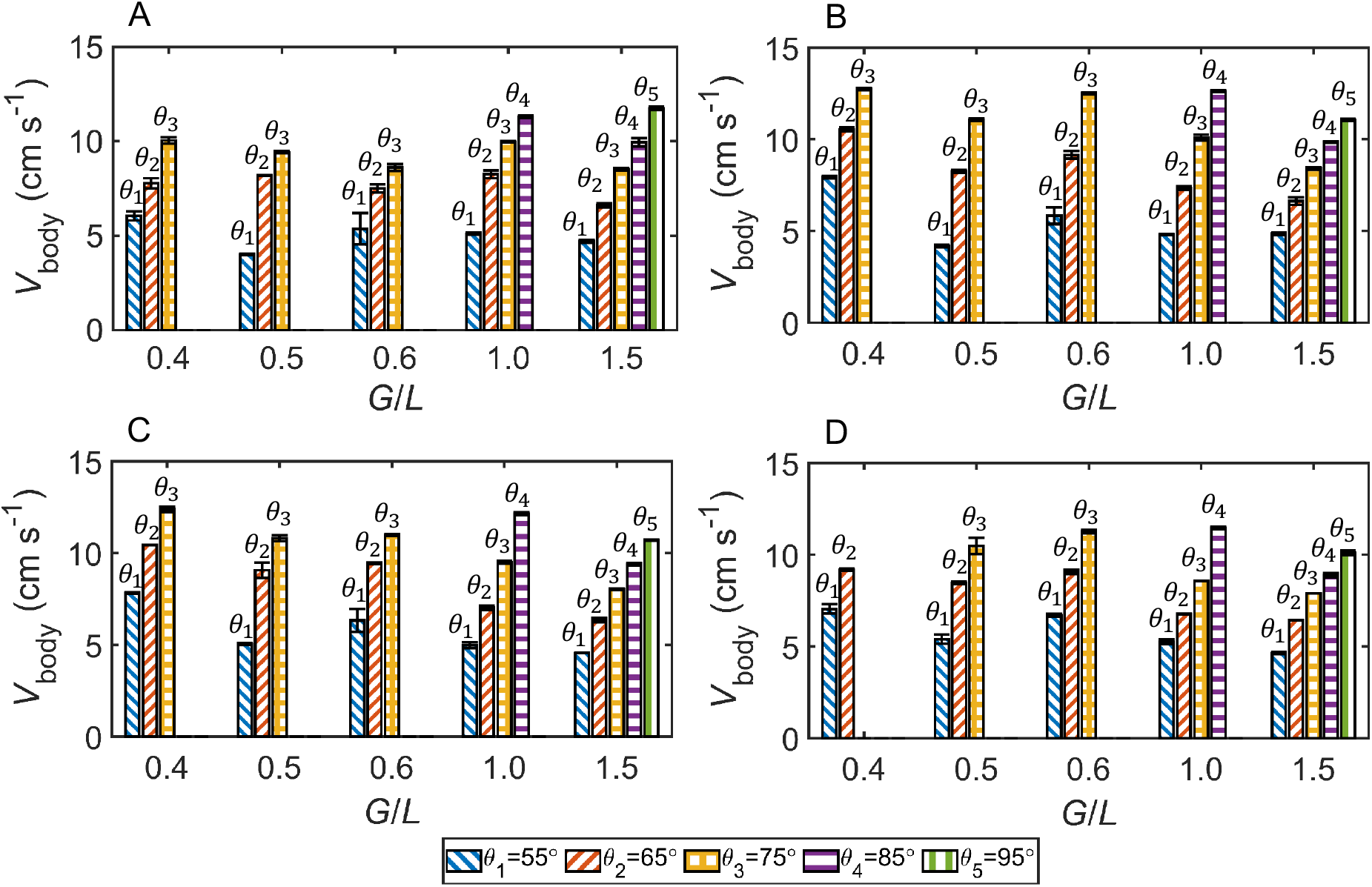
Steady swimming speed (*V*_body_) averaged over a stroke cycle. Bars represent varying stroke amplitude while groups of bars represent varying *G*/*L*. (A) phase lag, *ϕ* = 0%. (B) *ϕ* = 10%. (C) *ϕ* = 15%. (D) *ϕ* = 20%.

### 3.2 Flow field

To understand how the interaction of flows generated around individual paddles result in the large-scale wake of the paddling robot, we examined the velocity and normalized z-vorticity component (*ω_z_*/*ω*_*z*,max_) fields from PIV measurements (**Figure 3**). At *G*/*L*=0.5, *ϕ*=10%, *θ*=75° (**Figure 3A**) clockwise rotating tip vortices develop on each paddle during the power stroke (*t/T*=0 to 0.5), which are advected away from the body. By contrast, the counterclockwise rotating vortices generated by each paddle during the recovery stroke (*t*/*T*=0.5 to 1.0) are not advected away from the body, and instead merge to form one large region of vorticity rather than several smaller vortical structures. For *G*/*L*=1.5, *ϕ*=10%, *θ*=75° (**Figure 3B**), clockwise vortices are generated during power stroke of each individual paddle, similar to *G*/*L*=0.5 for the same *ϕ* and *θ*. However, unlike during the recovery stroke for *G*/*L*=0.5, the counterclockwise vortices generated during the recovery stroke with *G*/*L*=1.5 do not merge at either *θ*=75° (**Figure 3B**) or at *θ*=95° (**Figure 3C**). There is little variation seen in tip vortex formation and propagation when comparing *θ*=75° and *θ*=95° conditions at *G*/*L*=1.5, but both the velocity magnitude and the vortex strength increase with increasing *θ*. Additional flow fields for *G*/*L*=0.5 (with changing *ϕ*) and for *G*/*L*=1.0 are provided in supplementary material (**Figure S3**). Synchronous paddling (*ϕ*=0%) at *G*/*L*=0.5 results in large clockwise vortices being formed that promote reverse flow toward the anterior (head) of the model at the end of power stroke (**Figure S3,A**). At the end of power stroke, increasing phase lag to 20% at *G*/*L*=0.5 and *θ*=75° (**Figure S3,B**) results in more downward motion of flow compared to *ϕ*=10% for *G*/*L*=0.5 and *θ*=75° (**Figure 3A**). At *ϕ*=10%, slightly more separation of counterclockwise roating vortices are seen at the end of recovery stroke when comparing *G*/*L*=0.5 (**Figure 3A**, *t*/*T*=1.0) and *G*/*L*=1.0 (**Figure S3,F**).

**Figure 3.**
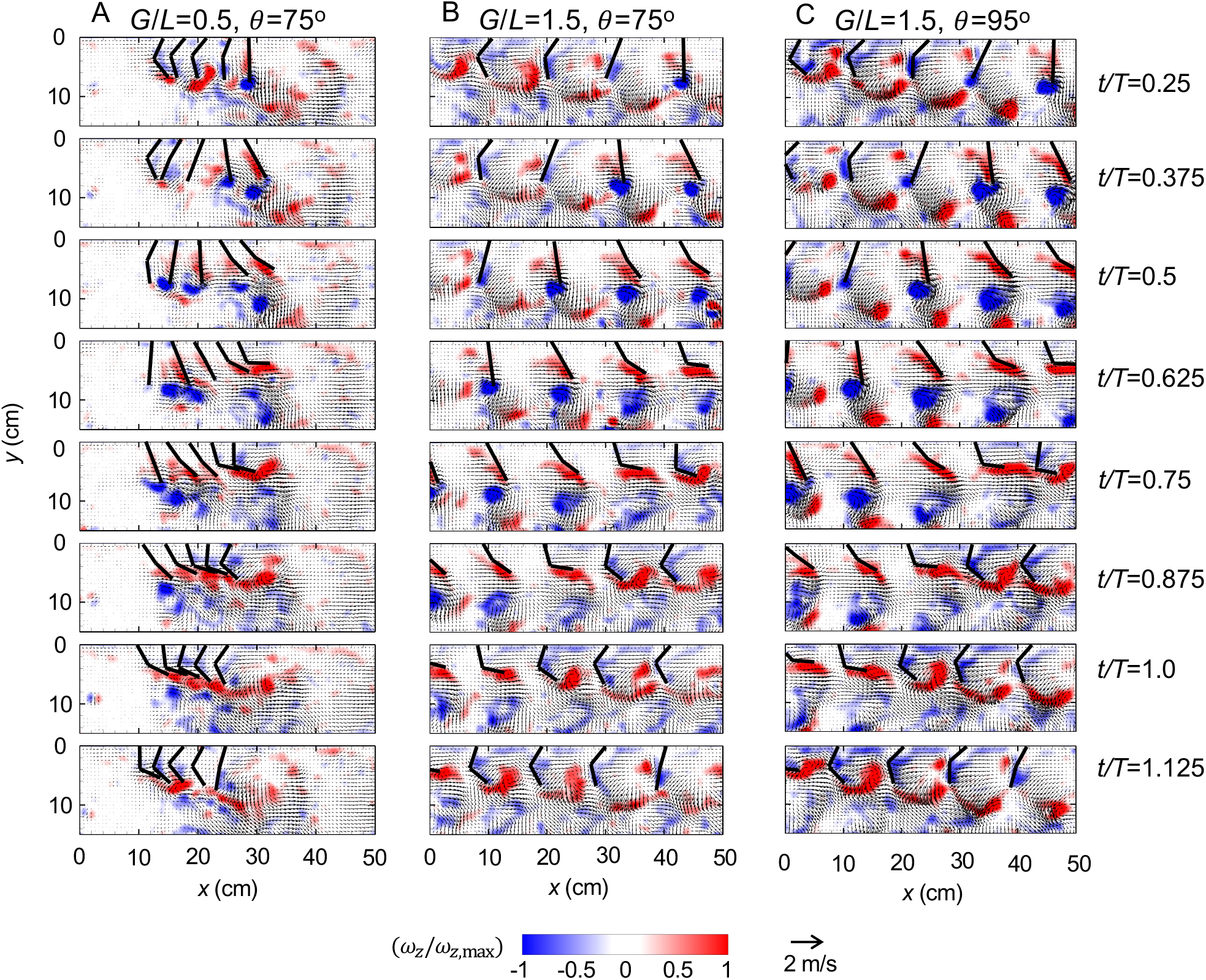
Vorticity contours overlaid with velocity fields for *ϕ*=10%. (A) *G*/*L*=0.5, *θ*=75°. (B) *G*/*L*=1.5, *θ*=75°. (C) *G*/*L*=1.5, *θ*=95°. When *G/L* is small, the wakes generated by individual paddles interact to form a large-scale downward jet. Non-dimensional times (*t/T*) indicated correspond to the P5 paddle (see **Figure 1**), with *t/T*=0 to 0.5 referring to duration of the power stroke, *t/T*=0.5 to 1.0 referring to duration of the recovery stroke. *t/T*=1 represents the end of the recovery stroke and the start of the subsequent power stroke.

### 3.3 Momentum flux

Horizontal and vertical momentum fluxes were calculated for the models with *G/L* = 0.5, 1.0 and 1.5. At *θ*=75°, changing phase lag has greater influence on the mean HMF (**Figure 4**), but also increases the cycle-to-cycle variation in HMF, as evidenced by the larger error bars. HMF is seen to be increasing along the entire body length for *G*/*L*=0.5, but not for *G*/*L*=1.0 or for *G*/*L*=1.5. For *G*/*L*=0.5, this augmentation of HMF with increasing horizontal distance signifies constructive interactions between the wakes of adjacent paddles. Multiple peaks are observed in the HMF data for *G*/*L*>0.5, located behind each pleopod, unlike the continuously increasing HMF values along the body length for *G*/*L*=0.5. This indicates that although fluid is advected along the body length, the wakes of the individual paddles are unable to interact to the same extent as at *G*/*L*=0.5, which allows some of the fluid momentum to be dissipated. The latter can in turn lower swimming speed for larger *G*/*L*, on account of reduction in useful (forward-directed) horizontal force generation. For each phase lag, *G*/*L*=0.5 has the highest HMF at the leeward (rear-facing) end of the model.

**Figure 4.**
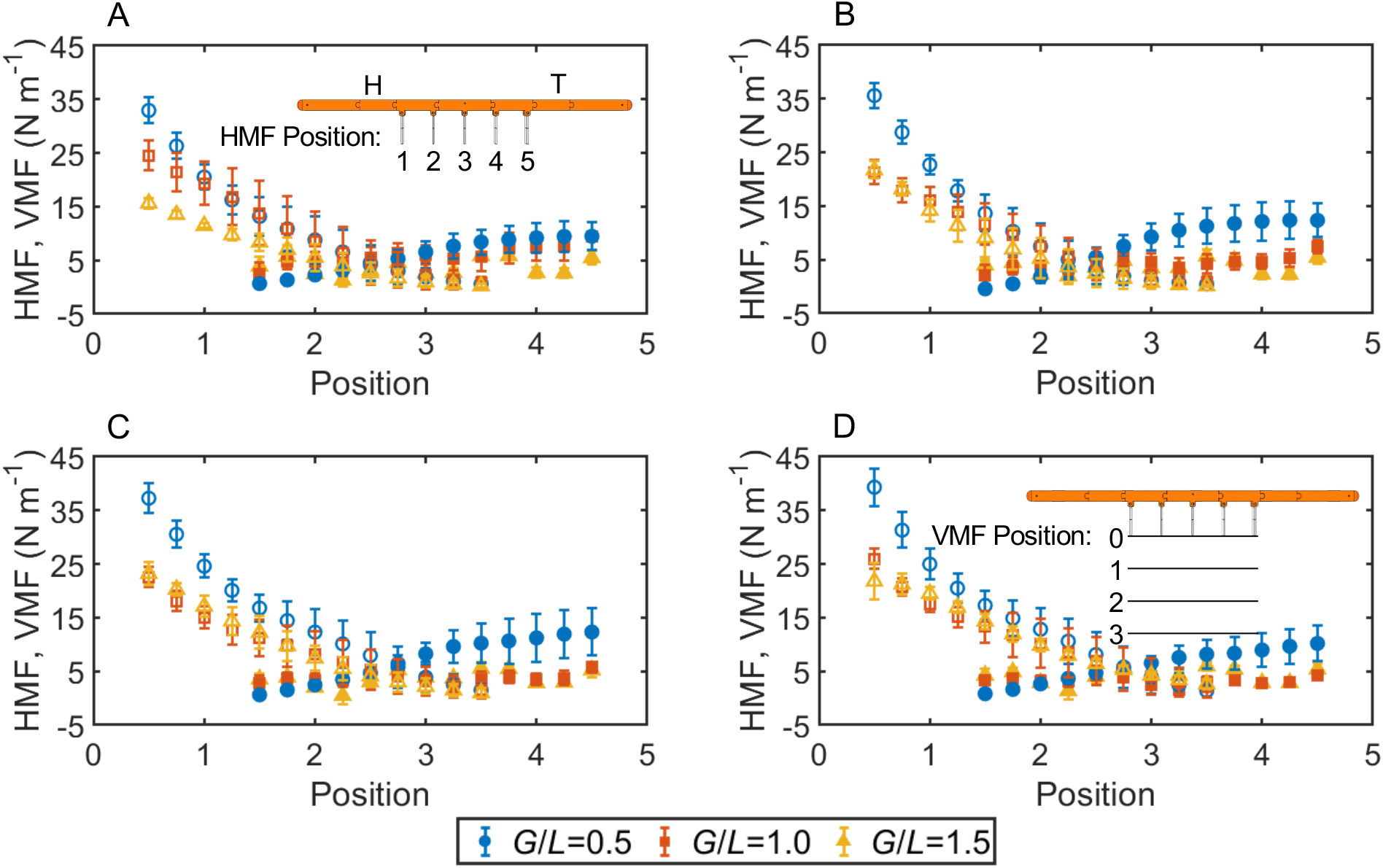
Vertical momentum flux (VMF, hollow markers) and horizontal momentum flux (HMF, filled markers) at *θ*=75°, measured at various depths below the body (for VMF) or positions along the body length (for HMF)., *x*-axis value for VMF indicates normalized distance below the body (in paddle lengths, see inset in part D)., *x*-axis value for HMF indicates the location along the body length, with 1 representing the location of P1, and 5 representing the location of P5 (see inset in part A). (A) *ϕ*=0%. (B) *ϕ*=10%. (C) *ϕ*=15%. (D) *θ*=20%.

In addition to HMF quantifying flow in the horizontal (thrust-generating) direction, VMF was used to quantify flow in the vertical or lift-generating direction (**Figure 4**). Over the first two paddle lengths below the body, VMF rapidly decreases due to viscous dissipation, and then dissipates slowly farther away from the body. Increasing *ϕ* results in increasing VMF, indicating that metachronal motion with non-zero phase lag is conducive in generating a strong vertical flow component. As with HMF, decreasing *G/L* from 1.5 to 0.5 results in increasing VMF. However, there is no effect of changing *G/L* from 1.0 to 1.5 on VMF for *ϕ*=10 to 20%, although *G*/*L*=1.0 does generate greater VMF than *G*/*L*=1.5 when *ϕ*=0%. Additional momentum flux data for *θ*=65° is presented in the supplementary material (**Figure S4**). While the trends for HMF variation with *ϕ* are unaltered when comparing *θ*=75°with *θ*=65°, VMF shows more separation between *G*/*L*=1.0 and *G*/*L*=1.5.

### 3.4 Wake angle

The overall momentum per unit depth was calculated within a box that covered the full length of the robotic model and extended 30 cm below the lower surface of the model (**Equation 6**). The horizontal and vertical components of momentum were used to determine the direction of the overall paddling wake using **Equation 7** and is shown in **Figure 5**. For a given value of *θ*, phase lag *ϕ* has far less effect on the angle of the paddling wake as compared to *G*/*L*. Likewise, for a given value of *ϕ*, stroke amplitude *θ* primarily affects only the magnitude of the momentum and not the angle of the jet. Changing the appendage spacing, on the other hand, results in significant changes in the wake angle. Larger *G/L* results in a more vertical wake that is not as conducive for motion in the horizontal direction, while smaller *G/L* results in a more horizontally directed jet that can augment swimming performance.

**Figure 5.**
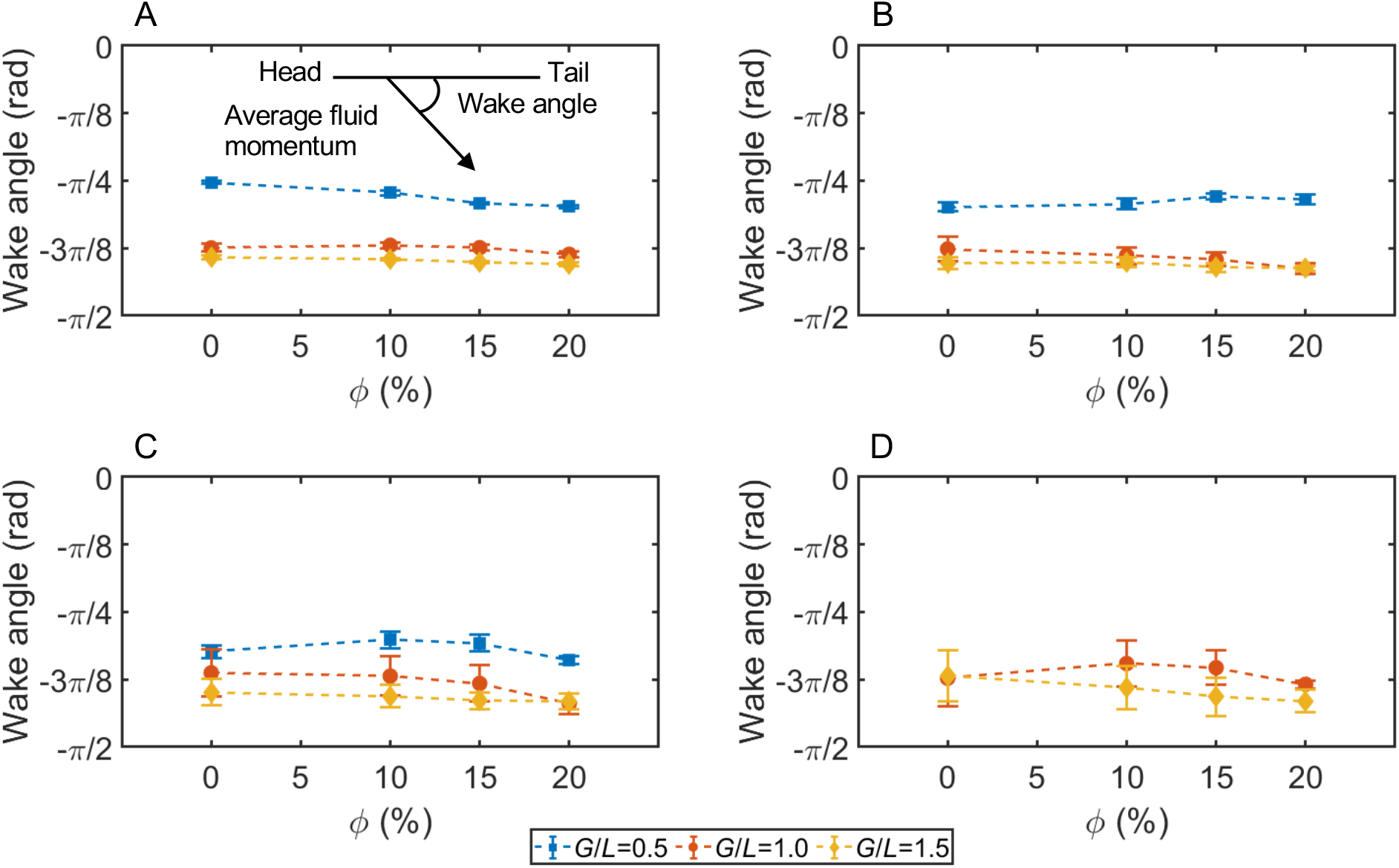
Angle of the overall wake generated by the paddling motion. An angle of 0 radians represents flow from head to tail of the robot, while an angle of −*π*/2 represents downward flow, perpendicular to the longitudinal axis of the body. (A) *θ*=55°. (B) *θ*=65°. (C) *θ*=75°. (D) *θ*=85°.

## 4 Discussion

In spite of numerous species-specific studies across a variety of crustaceans (Campos et al. 2012, Catton et al. 2011, Lenz et al. 2015, Lim and DeMont 2009, Murphy et al. 2011, Murphy et al. 2013, Schabes and Hamner 1992, Van Duren and Videler 2003, Yen et al. 2003), the functional significance of the narrow morphological variation in appendage spacing relative to appendage length (*G*/*L*) among species that use metachronal paddling for locomotion is unclear. The biological variation in *G/L* could serve a specific locomotor purpose, or could simply be a function of overall body morphology and energetics, since small organisms with long legs (such as copepods) simply cannot space their appendages as far apart as larger organisms can (such as stomatopods). This study examined the fluid dynamic effects of varying *G/L* to determine its effects on the metachronal wake and swimming performance. We hypothesized that for a given *Re* and stroke amplitude, hydrodynamic interactions promoted by metachronal stroke kinematics with small *G/L* can increase forward swimming speed. An implication of this statement is that when adjacent paddles are sufficiently far apart, there should be no hydrodynamic interactions and the swimming speed should be independent of both *ϕ* and *G*/*L*.

Using a dynamically scaled robotic paddling model with simplified geometry, we varied *G/L* and stroke kinematics (stroke amplitude *θ* and phase lag, *ϕ*) to determine how these variables affected swimming performance. Increasing *θ* resulted in higher swimming speeds for all conditions, and metachronal stroking at non-zero *ϕ* with *G/L* in the biological range produced faster swimming speeds and greater momentum fluxes than synchronous stroking. This is consistent with previous studies using closely spaced paddles (Alben et al. 2010, Zhang et al. 2014, Ford et al. 2021). To determine the *G/L* at which swimming performance becomes independent of these parameters, we examined models with *G*/*L* larger than biologically observed range of 0.2 to 0.65 (Murphy et al. 2011). For *G*/*L*=1.0 and *θ*=55°, and for *G*/*L*=1.5 and *θ* ≤ 65°, swimming speed was independent of *ϕ*. For larger *θ*, it was found that changing *ϕ* did indeed affect the swimming speed of the robot, but to a noticeably lesser extent than varying *ϕ* at *G*/*L*=0.5. Additionally, for larger *G*/*L*, larger *θ* was required to achieve the same swimming speed as for smaller *G/L* with smaller *θ*. Our results confirm that there is a minimum appendage spacing and tip velocity for the wakes of individual paddles to constructively interact so as to augment swimming speed. The distances between the hinges and the tips of adjacent paddles could serve as an indicator of inter-appendage hydrodynamic interactions. These hinge-to-hinge and tip-to-tip distances depend on *θ, ϕ, G/L* and hinge angle *β*, and are presented for *ϕ*=10% and *θ*=75° in the supplementary material (**Figure S5**).

The existing knowledge in the literature is that metachronal paddling, as opposed to synchronous paddling, is beneficial for swimming performance (Alben et al. 2010, Zhang et al. 2014, Ford et al. 2021). The novel contribution of this work is the finding that while metachronal motion augments swimming performance for closely spaced appendages (*G*/*L* < 1), moderately spaced appendages (1.0 ≤*G/L* ≤ 1.5) can benefit from metachronal motion only when the stroke amplitude is large. This finding can help inform the understanding of crustacean morphology vis-à-vis swimming performance, and can also be useful toward the engineering design of bio-inspired underwater vehicles.

### 4.1 Physical mechanisms

Interactions between the tip vortices of adjacent paddles during the power and recovery strokes seem to contribute to the swimming performance when multiple paddles move in a coordinated fashion. From the PIV data shown in **Figure 3** (and **Figure S3** in supplementary material) we can clearly see how the tip vortices in the near-wake of individual paddles with *G*/*L*=0.5 interact to form a coherent large-scale wake at *θ*=75°, such that *ϕ* has a noticeable effect on the swimming speed. Through much of the stroke at *G*/*L*=0.5, the tip vortices of each paddle are nearly indistinguishable in the large-scale wake of the paddling system. This is in marked contrast to the near-wake of the paddles when *G*/*L*=1.5 with *θ*=75° and with *θ*=95°, where *ϕ* shows minimal influence on the swimming speed. The hydrodynamic interactions at larger *G/L* are limited primarily to the counterclockwise vortices generated during the recovery stroke interacting with the leading edge of the immediately downstream (posterior) paddle.

The large-scale wake generated by the forward-swimming paddling system used in this study with *G*/*L*=0.5 is similar to the flow generated by a tethered 2-paddle system reported in our previous paper (Ford et al. 2019), where interactions between the counterrotating vortices generated during the power and recovery strokes aids in the generation of a primarily horizontal wake. While these time-dependent interactions happen on the level of an individual paddle for *G*/*L*=1.0 and 1.5, the wakes of the individual paddles do not merge to generate a large-scale wake. The interactions between the tip vortices generated by the power stroke with the merged vortices generated during the recovery stroke appears to be the fluid dynamic mechanism from which closely spaced appendages derive their thrust augmentation. This can explain why *G*/*L*=0.5 has a more horizontally angled wake (**Figure 5**) than the larger G/L with the same *θ* and *ϕ*, because the large-scale vortex interactions promoted by closely spaced appendages tailor the flow to move in a more horizontal direction.

### 4.2 Additional considerations

For a given *G*/*L*, stroke amplitude *θ* appears to be the strongest predictor of swimming speed. Phase lag *ϕ* influences swimming performance for smaller *G*/*L*. In order to avoid collisions between neighboring paddles, closely spaced paddles that move in a metachronal pattern (rather than a synchronous or nearly-synchronous pattern) are limited in the maximum *θ* that they can achieve when all paddles are held to a vertical mean angle (the angular paddle kinematics used in this study are mathematically represented by a sine wave with amplitude *θ* and mean value of 90°). There are a number of structural and kinematic innovations that could allow organisms to achieve greater stroke amplitudes while still using a series of closely spaced appendages for locomotion. Some of these innovations include: varying the mean paddle angle along the body (Murphy et al. 2011); using a different stroke plane for the power and recovery strokes (Schabes and Hamner 1992); separating the swimming stroke into a metachronal power stroke with synchronous recovery (Campos et al. 2012, Alexander 1988, Kiørboe et al. 2010); and using flexible appendages to reduce the risk of damage from collisions (Colin et al. 2020). In addition to reducing the risk of damage, flexible appendages have been shown to further augment force generation in flying and swimming with flapping appendages (Daniel 1988) and have been hypothesized to contribute to thrust generation in paddling arthropods (Colin et al. 2020). Individually or collectively, these innovations could contribute to swimming performance by allowing for increased stroke amplitude as well as by potentially affecting the wake development. These will need to be investigated individually in order to determine the unique contributions of each innovation to the metachronal swimming strategy.

## Acknowledgements

The authors would like to thank the following personnel at Oklahoma State University: Tyler Blackshare for his assistance with the design and manufacturing of parts; C. Tanner Price and Nicole Hackler for their assistance in acquiring high-speed videos of robot motion.

## Data Availability Statement

The data underlying this article are available in the article and in its online supplementary material.

