## Supplementary Figures S1 to S4 for "Closer appendage spacing augments metachronal swimming speed by promoting tip vortex interactions"

### SUPPLEMENTARY MATERIAL

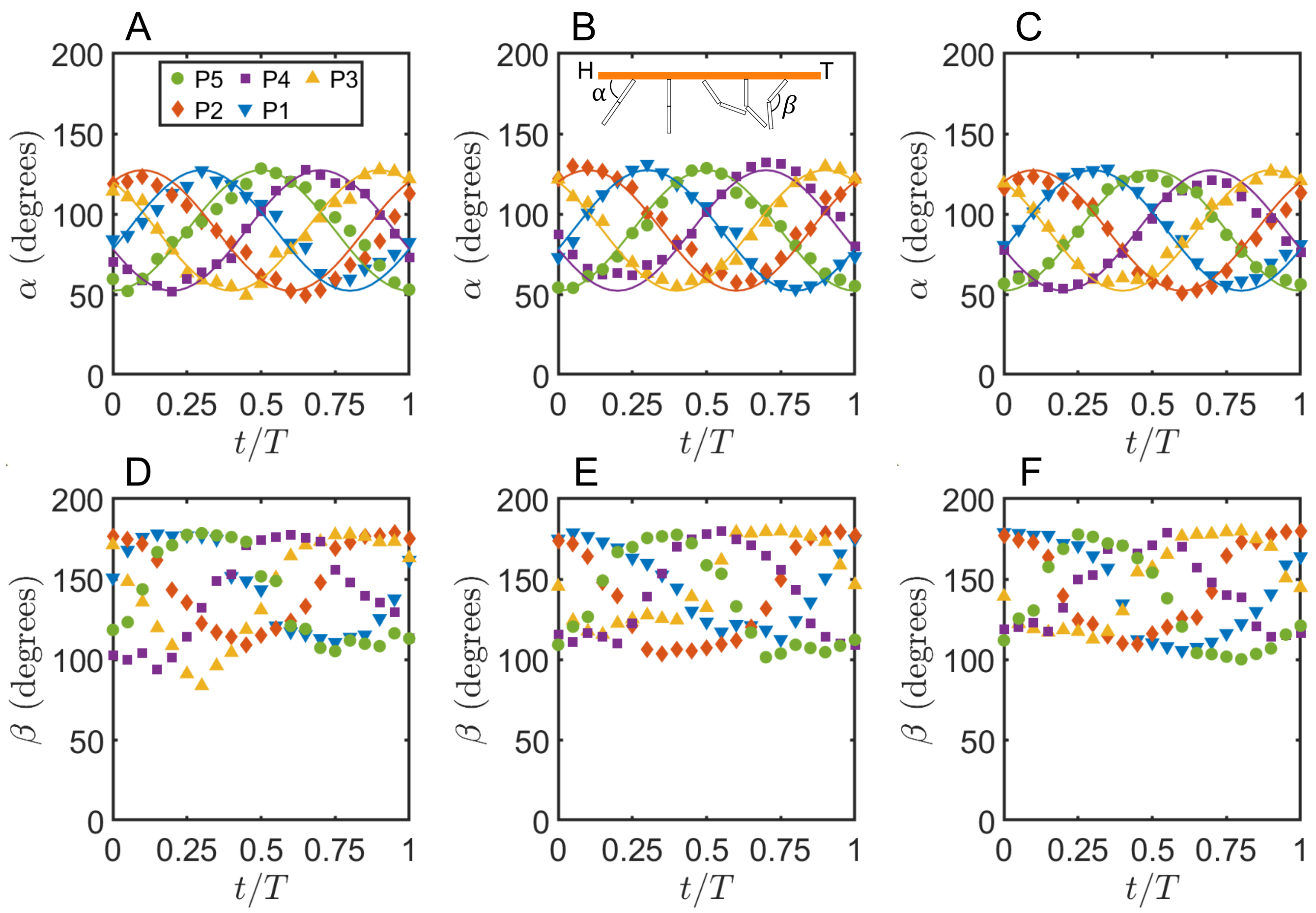


**Figure S1.** Tracked and prescribed kinematics for paddle motion with $\phi$ =10% and $\theta$ =75º. Definitions for appendage angle ($\alpha$) and hinge angle ($\beta$) are shown in the top of part (B). (A-C) $\alpha$ prescribed (solid lines) and tracked (markers) for models with (A) *G*/*L*=0.5, (B) *G*/*L*=1.0, and (C) *G*/*L*=1.5. (D-F) passive hinge angle $\beta$ tracked for each model (D) *G*/*L*=0.5, (E) *G*/*L*=1.0, and (F) *G*/*L*=1.5. $t/\tau$ is defined relative to the posterior paddle P5 (see **Figure 1** for paddle notations).

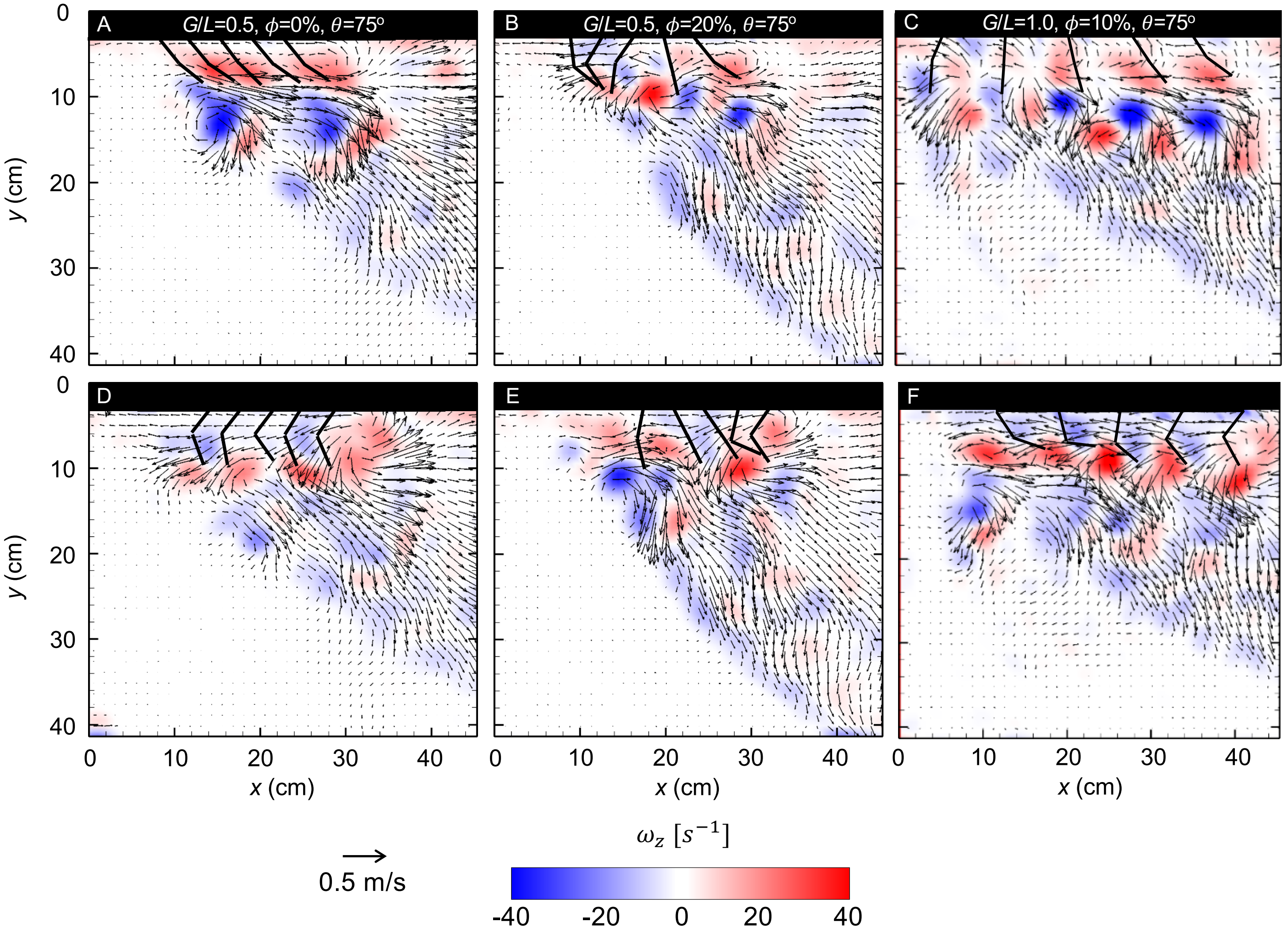


**Figure S2**. Velocity fields overlaid on vorticity contours for $G/L$=0.5, $\phi$=0%, $\theta$=75° (A&D), $G/L$=0.5, $\phi$=20%, $\theta$=75° (B&E), and $G/L$=1.0, $\phi$=10%, $\theta$=75° (C&F). (A-C) End of power stroke. (D-F) End of recovery stroke. Stroke instances are based on position of the posterior paddle P5 (see **Figure 1** for paddle notations). For $G/L$=0.5, the wakes generated by individual paddles are indistinguishable from each other in the large-scale wake. The synchronous motion gives a pulsed wake directed in a more horizontal direction than in the $\phi$=10% case shown in **Figure 3**, and in the $\phi$=20% case shown here. Increasing the limb spacing allows for increasing stroke amplitude while avoiding collisions between neighboring limbs. For $G/L$=1.0, counter rotating vortex pairs can be clearly seen near the tip of each paddle as it completes the power stroke.

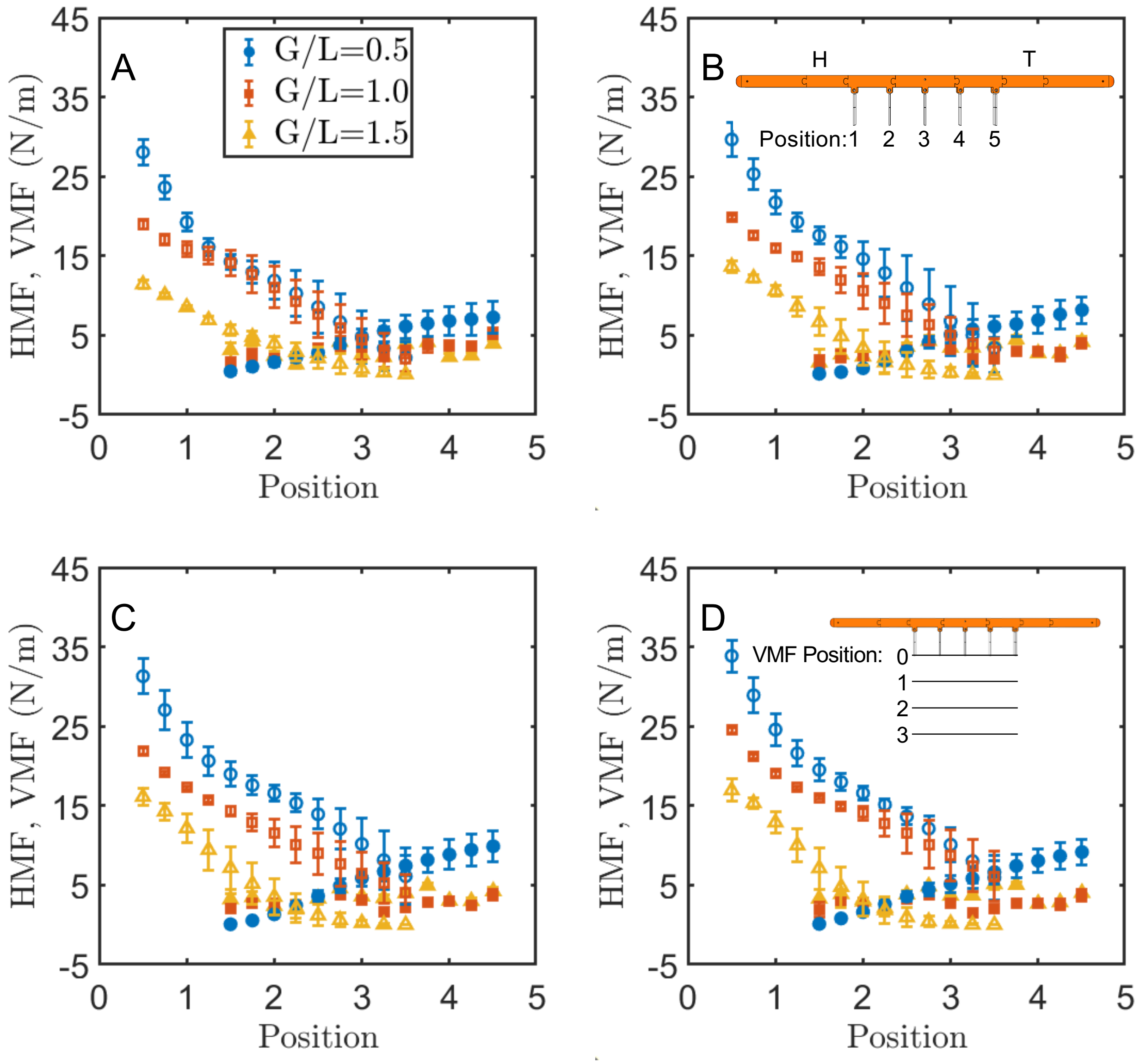


**Figure S3**. Vertical (dashed lines) and horizontal (solid lines) momentum flux at $\theta$=65º, measured at various depths below the body (for VMF) or positions along the body length (for HMF). $x$-axis value for VMF indicates normalized distance below the body (in paddle lengths, see inset in part D). $x$-axis value for HMF indicates the physical location along the body length, with 1 representing the location of P1, and 5 representing the location of P5 (see inset in part B). (A) $\phi$ =0%. (B) $\phi$ =10%. (C) $\phi$ =15%. (D) $\phi$ =20%.

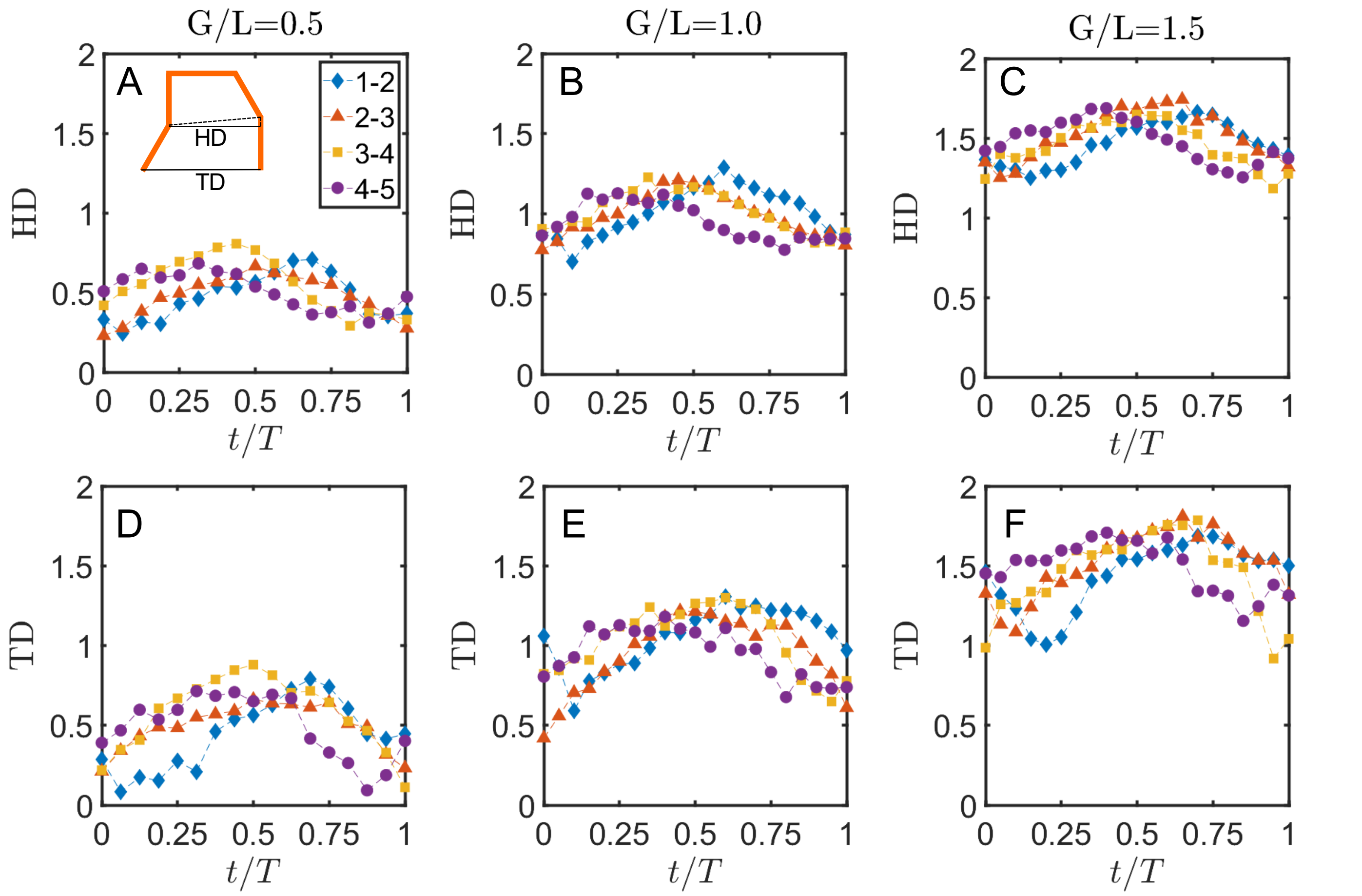


**Figure S4.** Time-histories of the distances between hinges and tips of neighboring paddles, normalized by paddle length. (A) hinge-distances for *G*/*L*=0.5, $\phi$=10%, $\theta$=75º. (B) hinge-distances for *G*/*L*=1.0, $\phi$=10%, $\theta$=75º. (C) hinge-distances for *G*/*L*=1.5, $\phi$=10%, $\theta$=75º. (D) tip-distances for *G*/*L*=0.5, $\phi$=10%, $\theta$=75º. (E) tip-distances for *G*/*L*=1.0, $\phi$=10%, $\theta$=75º. (F) tip-distances for *G*/*L*=1.5, $\phi$=10%, $\theta$=75º.
